# Multiple interfacial hydration of dihydro-sphingomyelin bilayer reported by the Laurdan fluorescence

**DOI:** 10.1101/391128

**Authors:** N. Watanabe (N. W.), Y. Goto (Y. G), K. Suga (K. S.), T. Nyholm (T. N.), J. P. Slotte (J. P. S.), H. Umakoshi (H. U.)

## Abstract

The hydration properties of the lipid bilayer interface are important for determining membrane characteristics. The hydration properties of different lipid bilayer species were evaluated using the solvent sensitive fluorescence probe, 6-lauroyl-2-dimethylamino naphthalene (Laurdan). Sphingolipids, D-*erythro*-N-palmitoyl-sphingosylphosphorylcholine (PSM) and D-*erythro*-N-palmitoyl-dihydrosphingomyelin (DHPSM) showed specific, interfacial hydration properties stemming from their intra- and intermolecular hydrogen bonds. As control, the bilayers of glycerophospholipids, such as 1-palmitoyl-2-palmitoyl-*sn*-glycero-3-phosphocholine (DPPC) and 1-oleoyl-2-oleoyl-*sn*-glycero-3-phosphocholine (DOPC), were also evaluated. The fluorescence properties of Laurdan in sphingolipids indicated multiple excited states according to the results obtained from the emission spectra, fluorescence anisotropy, and the center of mass spectra during the decay time. Deconvolution of the Laurdan emission spectra into four components enabled us to identify the variety of hydration and the configurational states derived from intermolecular hydrogen bonding in sphingolipids. Particularly, the Laurdan in DHPSM revealed more hydrated properties compared to the case in PSM, even though DHPSM has a higher T_m_ than PSM. Since DHPSM forms hydrogen bonds with water molecules (in 2NH configurational functional groups) and the different flexibility among the head groups compared with PSM, which could modulate space to retain a high amount of water molecules. The careful analysis of Laurdan such as the deconvolution of emission spectra into four components performed in this study gives the important view for understanding the membrane hydration property.

## Introduction

Plasma membrane bilayers play a fundamental role of regulating cellular function by delineating a membrane wall which divides the intracellular and the extracellular spaces. In addition, each role of the membrane, such as encapsulation, localization of proteins, and molecular permeation across the membrane, can depend on the compositions of their membrane proteins and lipid species (1). In particular, lipid rafts have been attracting interest of researchers for many years because they mediate important cellular functions via interaction with biomolecules (2–5). Once lipid rafts are formed in the bilayer, they could be stabilized through hydrogen bond networks between lipid molecules (3, 6). Sphingolipids are the major lipid species known to associate with raft domain stability (7). The intermolecular hydrogen bond can be facilitated through their hydrogen bonding acceptor (C=O, P=O, 3OH, or 2NH) and donor moieties (3OH or 2NH) (6, 8). Furthermore, most of sphingolipids have long, saturated acyl chain that allows them to be packed tightly (9). Even with a subtle configurational difference of saturated or unsaturated, the physicochemical property, such as the phase transition temperature (*T*_m_) of sphingolipids in bilayer, can be varied. For example, D-*erythro*-N-palmitoyl-sphingosylphosphorylcholine (PSM), having a saturated acyl chain and an unsaturated long-chain base, shows the *T*_m_ of 41.2 °C in bilayer (10). D-*erythro*-N-palmitoyl-dihydrosphingomyelin (DHPSM), having a saturated long-chain base, showed the *T*_m_ of 47.7 °C in bilayer (11–13). The former and later sphingolipids are found as a predominant species in most of plasma cell membranes and in human lens cell membranes, respectively. The hydrogen bonds between lipids makes the phase state of the membrane rigid and ordered, especially in the presence of cholesterol, which can lead to important raft functions, such as localization and transportation of proteins during signal transduction.

Integration of external (or internal) biomolecules into the membrane surface is promoted by several factors: the electrostatic and hydrophobic interactions between those molecules and the membranes are usually dominant (14–16). Furthermore, the water molecules at the interface of lipid membrane can control the activities of biomolecules by modulating surface pressure, coordination of hydrogen bonding, and surface charge state (17, 18). In lipid raft, the membrane interface could be maintained as dehydrated state, because the intermolecular hydrogen bonding between sphingolipids can exclude the hydration waters. Therefore, it is considered that the hydration property at the membrane interface could regulate raft function which is able to integrate molecules and promote their activity. Many researchers have studied membrane hydration (18–21). It could be hypothesized that sphingomyelins may have specific hydration properties at the bilayer surface. However, little is known regarding the interfacial hydration state of local raft regions compared to other regions within the same membrane, because the observation of heterogenic hydration properties has not been established.

To evaluate membrane hydration states, the fluorescence probe 6-lauroyl-2-dimethylamino naphthalene (Laurdan) has been widely used (22–24). The fluorescence emission of Laurdan is sensitive to the surrounding solvent environment. In the lipid bilayer, the amphipathic structure of Laurdan localizes it at the hydrophobic-hydrophilic interface region. Thus, the obtained information are relevant to the microscopic interfacial properties of the lipid bilayer (24, 25). By excitation, the fluorophore moiety has a large dipole moment (25, 26). Laurdan can take multiple excitation energy states. The highest excitation energy state can be relaxed by surrounding solvent molecules, which results in a complex emission property, such as blue shift or long lifetime (24, 27, 28). The predominant emission peaks of the Laurdan in lipid bilayer have been previously identified, one can be observed in the gel phase membrane with the *λ*_em_ of 440 nm, and the other can be observed in the liquid-crystalline phase membrane with the *λ*_em_ of 490 nm) (22).

By utilizing these emission properties of Laurdan in lipid bilayer, an analytical method for the microscopic polarity of the membrane has been developed: considering two-state assumption (gel-phase or liquid phase), the generalized polarization (GP) value can be used to assess membrane hydration during phase transition (22). The relaxation state of Laurdan can be measured through deconvolution of the emission spectrum, measurement of fluorescence lifetime, and the transition of the center of mass spectra in time-resolved fluorescence analysis, which can also be used to assess membrane hydration (25, 29–31). Despite these methodologies, further investigation has been required to evaluate dynamic hydration behaviors at the lipid bilayer surfaces, since there are many factors that would affect the heterogeneity of Laurdan relaxation, e.g., hydrogen bonding, lateral lipid density, lateral heterogeneity, and so on (32).

Once again, the GP method can be widely applied to investigate the hydration properties of the membrane, while subtle differences in the Laurdan emission peaks found in sphingolipid bilayer (33), have not been discussed enough. In this study, we evaluated the interfacial hydration properties of bilayers made from different lipid species to determine whether the different configurations, including backbone structure or degree of saturation, would affect the surface hydration properties of bilayer. Specifically, we focused on how the hydrogen-bonding donor and acceptor groups in PSM and DHPSM affected the interfacial hydration. To evaluate the hydration properties of bilayer membrane, the steady state measurements of excitation spectra, emission spectra, and anisotropy of Laurdan were carefully performed, together with the time-resolved emission analysis. Herein, we propose the deconvolution of obtained Laurdan fluorescence emission spectra into four components. Because the emission properties of Laurdan depend on its surroundings, the differences of microscopic surroundings around Laurdan in each lipid bilayer system can be discussed as “hydration property” of the lipid bilayer surface.

## Materials and Methods

### Materials

All phospholipids (egg yolk SM, 1-palmitoyl-2-palmitoyl-*sn*-glycero-3-phosphocholine (DPPC), 1-oleoyl-2-oleoyl-*sn*-glycero-3-phosphocholine (DOPC), and 1-palmitoyl-2-oleoyl-*sn*-glycero-3-phosphocholine (POPC)) were purchased from Avanti Polar Lipids (Alabaster, AL, USA). PSM was purified from egg yolk SM as described previously (12). PSM was hydrogenated to yield DHPSM, as previously reported (9). The purity of sphingomyelins was confirmed by mass spectrometry. Laurdan was purchased from Sigma Aldrich (St. Louis, MO, USA). Other chemicals were obtained from Wako Pure Chemical Industries, Ltd. (Osaka, Japan), and used without further purification.

### Preparation of the lipid bilayer vesicles

Lipids and fluorescence probe were dissolved in methanol. Appropriate amounts were dispensed into a glass tubes, and the solvent was evaporated under a stream of nitrogen gas. The dry lipid films were hydrated in pure water at 65 °C (60 min) and sonicated in the sonicator bath, Bransonic 2510 (Branson Ultrasonics, CT, USA), for 5 min at 65 °C. Large unilamellar vesicles (LUVs, 100 nm diameter size) were prepared by extrusion (34). Lipid concentration was measured using the method reported by Rouser et al (35).

### Fluorescence measurements of Laurdan in steady state

All samples were prepared to adjust a final concentration of lipid and Laurdan to 100 μM and to 1 μM, respectively. The concentration of Laurdan was estimated from its absorbance, using the molecular extinction coefficient (ε = 2000 cm^−1^) (36). Steady state fluorescence measurements were performed with a QuantaMaster™ from Photon Technologies International (Lawrenceville, NJ, USA). The temperature of the sample cell was monitored and regulated by a Peltier system, and data acquisition was controlled using the Felix 32 software. Emission measurement was performed using excitation light at 360 nm, and emission spectra of Laurdan were collected from 400 to 540 nm.

Fluorescence anisotropy, *r*, was calculated from vertically (*I*_⊥_) and horizontally (*I*_║_) polarized emission intensity by the following equation;

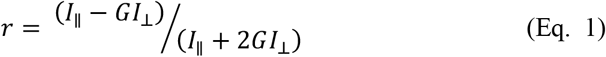

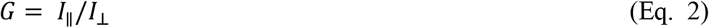

as defined in a previous report (37). The changes of anisotropy as a function of temperature were continuously measured from 20 °C to 70 °C, and the results were analyzed with the sigmoidal fitting curves, based on Boltzmann equation. The measurement was performed using excitation light at 360 nm, and the emission was measured at 490 nm.

The deconvolution of obtained emission spectra was carried out using the software Peakfit (Systat Software, Inc., San Jose, CA, USA). All fitting equations were operated with the lognormal amplitude function following a previous report (30). The equation for the fitting is;

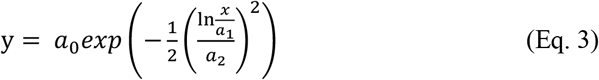

where *a*_0_ is the amplitude, *a*_1_ is the center (≠0) and *a*_2_ is the width (>0) of the fitting curve, respectively. From the fitting acquisition, the integrated area and its area ratio [%] were obtained; the integrated area was calculated from the integration of the obtained analytical curve.

### Time-resolved fluorescence measurements

Time-resolved fluorescence measurements were all performed on the FluoTime 200 instrument (PicoQuant, Rudower Chaussee, Berlin, Germany). The sample temperature was controlled by a Peltier system, and data acquisition was performed with the PicoHarp system. The samples were excited with a 378 nm diode laser and the emission was measured from 400 to 540 nm in 10 nm step (14 acquisitions). The time resolution was performed with 64 picoseconds. A decay curve was produced after150 seconds acquisition time. Data were analyzed by the FluoFit Pro software. After integration of all decay curves, the time-resolved emission spectra were extracted in every 1.6 nanoseconds after excitation (from 7.168 ns to 48.768 ns, in 28 spectra), and the center of spectral mass (*λ*) was calculated according to the previous report (38);

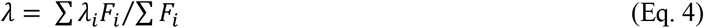

where *F_i_* is the fluorescence emitted at the emission wavelength *λ_i_*.

## Results

### Emission spectra of Laurdan in the steady state

The fluorescence emission spectra of Laurdan were measured in the presence of bilayers formed by DPPC, DOPC, PSM, and DHPSM. As shown in Figure 1, the Laurdan in DPPC (T_m_: 41°C) showed a sharp peak at around 440 nm, which was well-consistent with the reported Laurdan peak position of DPPC bilayer in gel phase state (22). In contrast, the Laurdan spectrum in DOPC bilayer showed a broader, red-shifted peak at approximately 490 nm, which can be derived from the Laurdan located in liquid crystalline phase. For the Laurdan molecules in PSM and DHPSM bilayers at 20 °C, the spectra showed broader peaks compared with that in DPPC. The phase transition temperatures of PSM and DHPSM in bilayer are 41.2 °C and 47.7 °C, respectively (10), suggesting that the phase states of PSM and DHPSM bilayers could resemble that of DPPC bilayer. In the SM bilayers, broader peaks, at 440 nm in PSM and at 460 nm in DHPSM, indicate that the Laurdan in SM bilayer would be in multiple hydration states. A red-shifted spectrum was observed in DHPSM bilayer compared to PSM, despite the lateral packing state of DHPSM and PSM molecules could be almost similar (39). This result suggests that DHPSM bilayer may be highly hydrated at the lipid bilayer interface where Laurdan existed. The Laurdan emission spectrum in DHPSM bilayer seems to be considerably similar to that in DOPC bilayer, suggesting this lipid can form a highly hydrated bilayer similar to DOPC bilayer.

**Figure 1.**
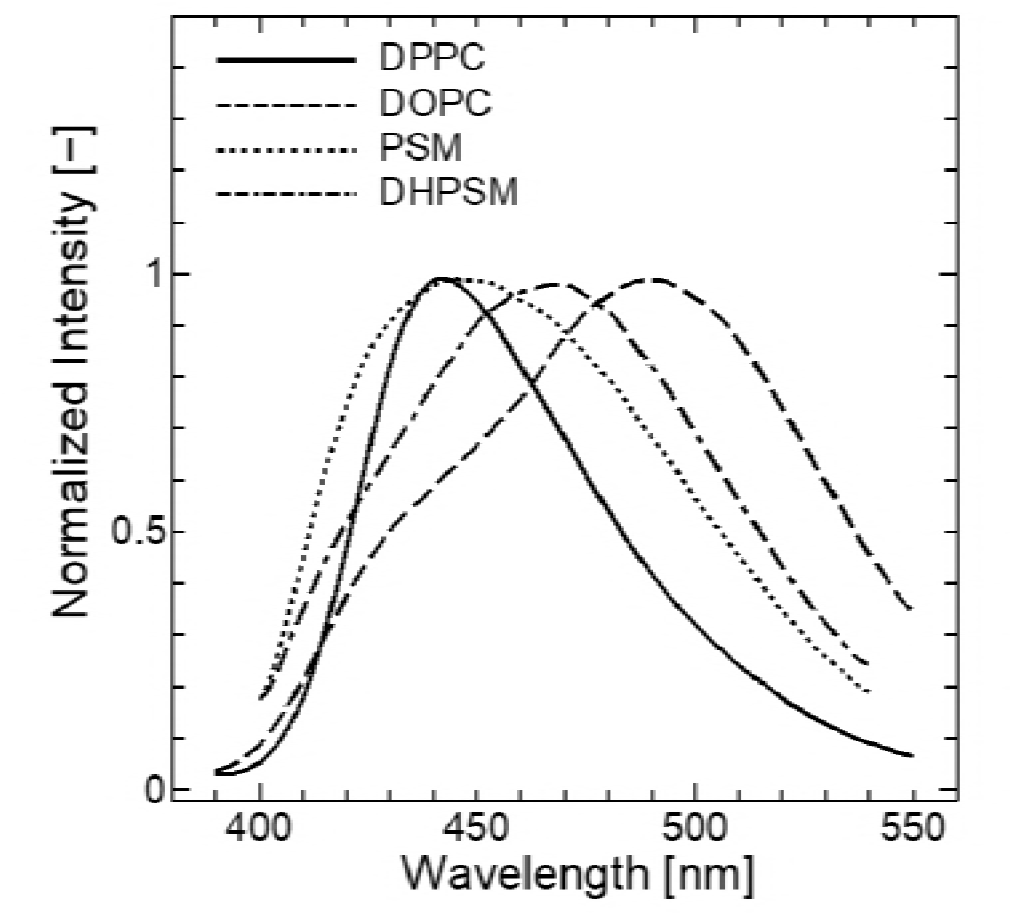
Fluorescence emission spectra of Laurdan in DPPC (solid), DOPC (dash), PSM (dot), and DHPSM (dot 2-dash) bilayers at 20°C. The excitation wavelength was 360 nm. All spectra were normalized with the strongest peak intensity. The reproducibility found from the different compositional binary system were also shown in Figure S1.

### Anisotropy of Laurdan

The rotational mobilities of the fluorescent molecules in bilayer were investigated by measuring their steady state anisotropy. The anisotropy value would decrease if the molecules were put into more disordered environment, wherein the molecular rotational mobility or freedom would increase: for example the phase transition from gel to liquid-crystalline phase. The anisotropy values of Laurdan in each lipid bilayer are shown in Figure 2. In DPPC bilayer, a drastic shift of anisotropy was observed at the temperature range around T_m_ (= 41°C). Since the DOPC bilayer is fluid and disordered state in this temperature range, there was only a slight response in the anisotropy of Laurdan to the temperature. PSM and DHPSM bilayers showed broader shifts of anisotropy values, as compared to the shift observed in DPPC bilayer systems. Moreover, At the temperature below T_m_, the Laurdan in DHPSM bilayer showed relatively smaller anisotropy values in comparison to those in DPPC bilayer in the gel phase, indicating that the Laurdan molecules might have high mobility at the DHPSM bilayer interface. The increase of dipole alignment of lipid head group was observed in low water content (40). This implies that a disordered lipid alignment can provide more hydrated water the bilayer interface, in such environment, a possibility of the contact between water molecules and located probe molecules could be increased. Considering gradual shifts of anisotropy values in SM bilayers, compared to glycerophospholipid species, it suggests the presence of more hydrated waters at the interface region, despite their tightly-packed state below Tm. In common to all samples, a large scattering in the anisotropy values was observed, this could be caused by a variety of the orientation of Laurdan in each lipid bilayer.

**Figure 2.**
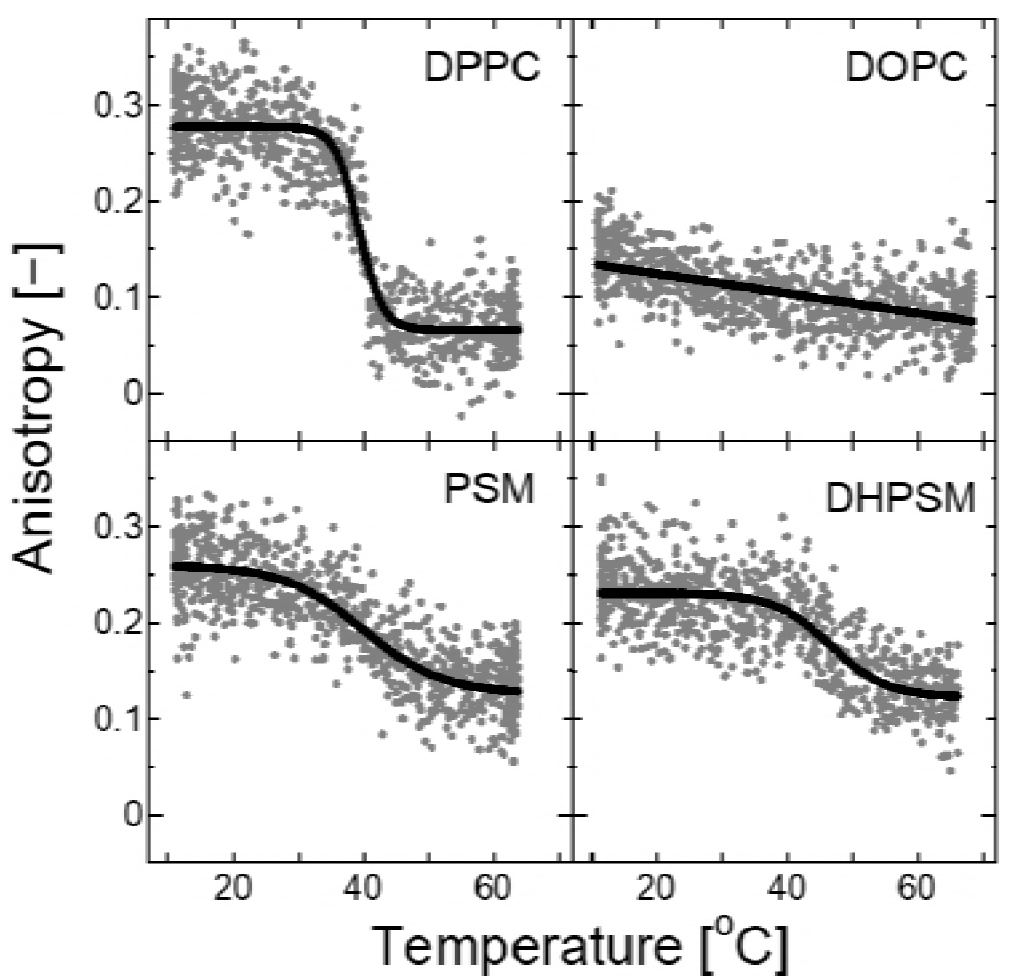
Anisotropy transitions as a function of the temperature in DPPC, DOPC, PSM, and DHPSM bilayers. The excitation wavelength was 360 nm, and the emission wavelength was at 490 nm. Anisotropy was calculated using the Eq. 1 and 2. The sigmoidal fitting curve was illustrated using the Boltzmann equation. The reproducibility found from the different compositional binary system are also shown in Figure S2.

### Center of mass of the time-resolved emission spectra

Time-resolved fluorescence analysis is a powerful tool to interpret molecular dynamics of fluorescence probes. The fluorescence emission decay can be calculated and the center of mass of the spectrum can be obtained from the integration of the whole emission range. The center of mass of the time-resolved emission spectra of Laurdan was calculated for DPPC, DOPC, PSM, and DHPSM bilayers (Figure 3). The significant red-shift in DOPC bilayer was observed, whereas the value was only slightly shifted in DPPC bilayer. These tendencies are in agreement with previous findings (29). For the sphingolipids, the center of mass of DHPSM bilayer was significantly red-shifted, whereas PSM bilayer showed a transition similar to DPPC bilayer. A unique feature of sphingolipids was found as a function of fluorescence decay time. In SM bilayers, the center of mass shifted to large wavelength in shorter decay time, indicating a quick response of Laurdan to the solvent relaxation from the surrounding hydrated waters. This observation suggests that there are specific molecular interactions among sphingolipids that enable the probe to be stabilized and relaxed in a short time.

**Figure 3.**
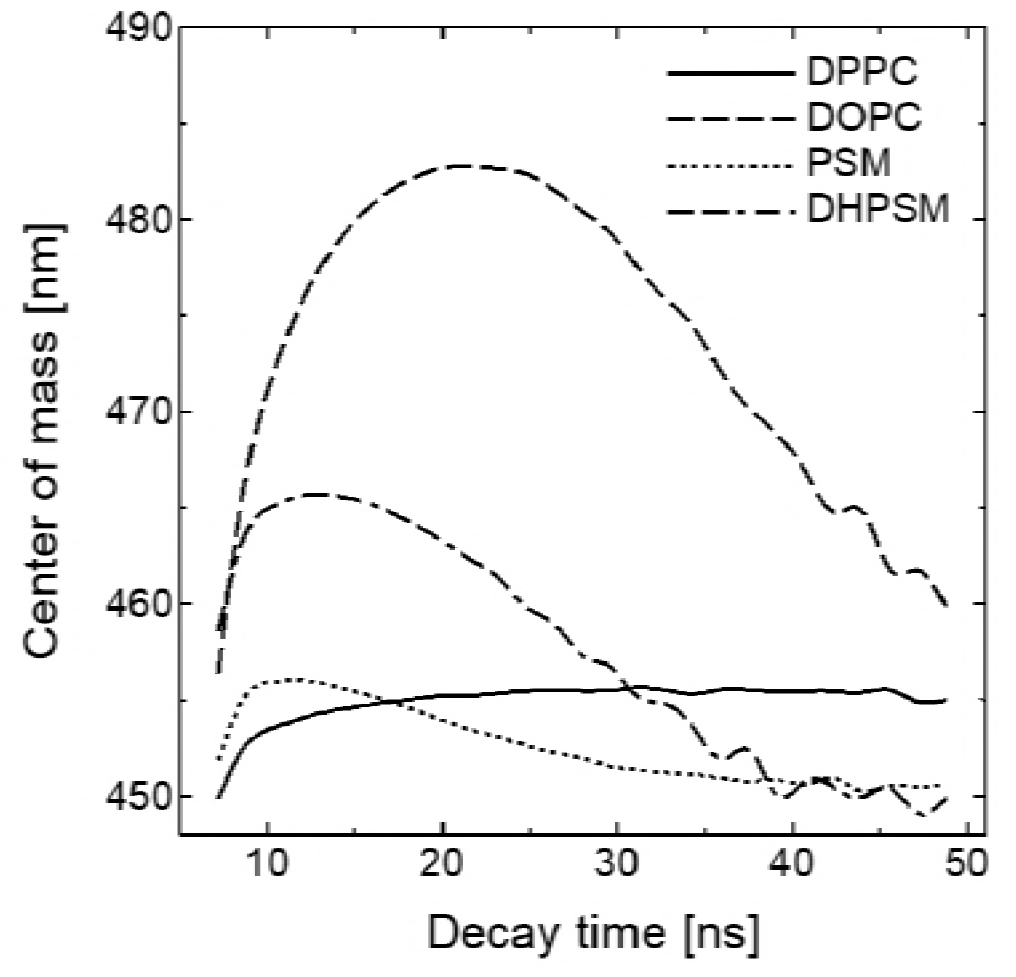
The center of mass of the time-resolved spectra of Laurdan for each bilayer depicted as DPPC (solid), DOPC (dash), PSM (dot), and DHPSM (dot 2-dash) at 20°C. The excitation wavelength for the diode laser was 378 nm. The value of the center of mass was calculated using Eq. 4. Each data points were extracted every 7.168 ns. The reproducibility found from the different compositional binary system are also shown in Figure S3.

### Excitation spectrum of Laurdan in DOPC and SMs

The intermolecular interactions between Laurdan and lipids were investigated by measuring the excitation spectra by Parasassi *et al*. They investigated that Laurdan can be stabilized via the interaction with the ester groups of the glycerophospholipid by observing the relative intensities of two excitation bands (24). From the comparison with glycerophospholipid, they suggested the interaction of Laurdan with sphingolipid can be weaker because the inter lipid hydrogen bonds among sphingolipids are strong so that Laurdan couldn’t interact with lipids. In Figure 4, the excitation spectra for Laurdan in PSM and DHPSM bilayers showed red-shifts. The red-shifted excitation spectra of sphingolipid were also observed in their results (24). Generally, the excitation spectrum is symmetric with that of the emission spectrum known as Franck-Condon principle (41). Moreover, some environmental factors, such as intermolecular interaction or temperature, could stabilize probe molecules and enable them to appear in furthermore relaxed state, resulting in a lower excited state (long wavelength range) (37). Based on the theoretical backgrounds, the red-shifted excitation spectrum of Laurdan in SM bilayers could be explained due to the configurational property such as intermolecular hydrogen bonds, resulting in the blue-shifted emission spectrum shown in Figure 1.

**Figure 4.**
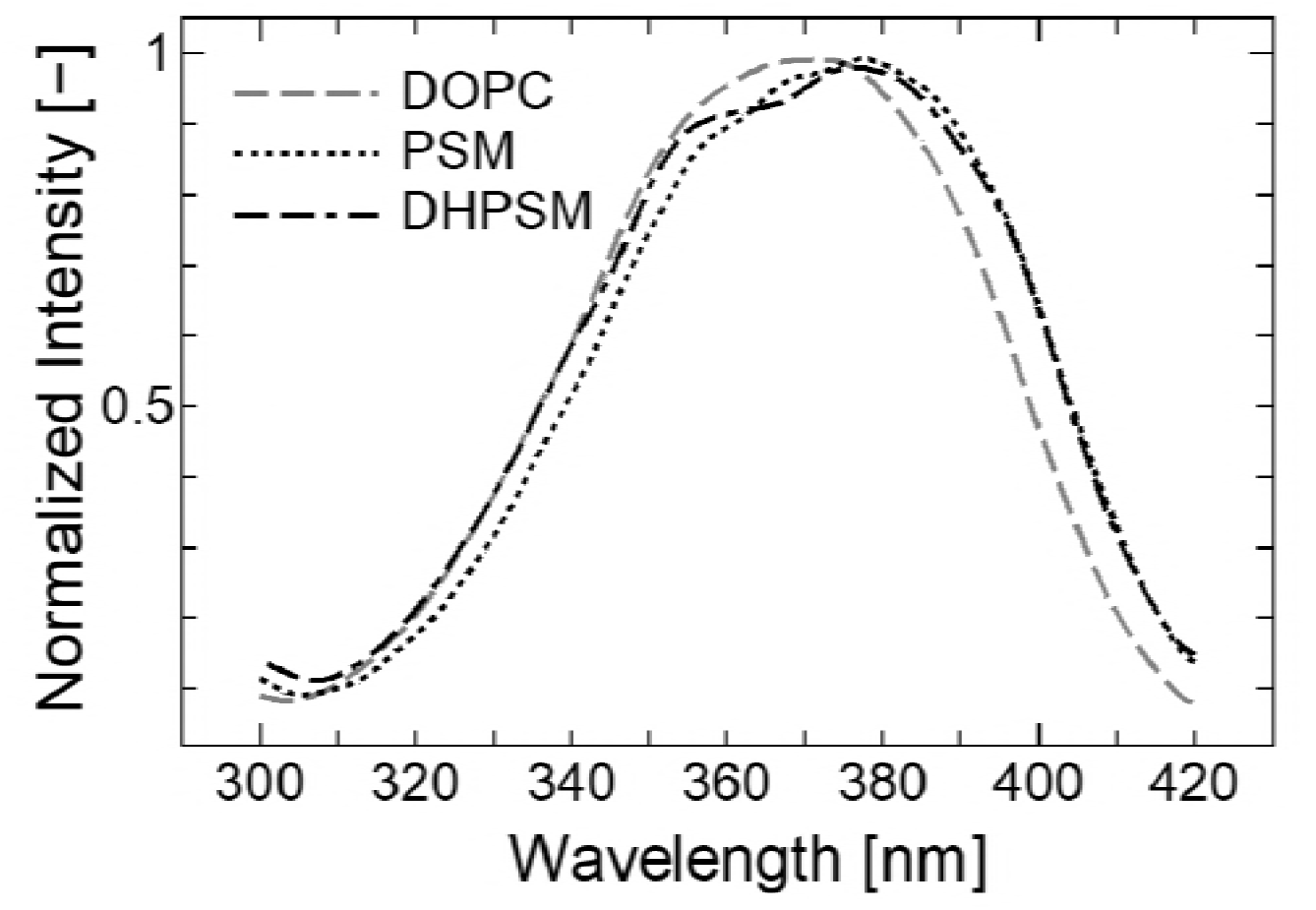
Laurdan excitation spectra of DOPC (dash), PSM (dot) and DHPSM (dot 2-dash) bilayers at emission wavelength (λ_em_ = 435 nm) at 20°C. All spectra were normalized with the strongest peak intensity. The reproducibility found from the different compositional binary system are also shown in Figure S4.

## Discussion

### Interpretation of hydration states in PC and SM bilayers using the fluorescence properties of Laurdan

Analysis of the Laurdan emission spectra is a useful tool to characterize membrane bilayer interfaces. Some spectral features of the fluorescence emission from representative lipid bilayers are known. For example, the gel-like membrane phase is indicated by a narrower significant blue-shifted emission peak while a broader red-shifted peak indicates a relatively disordered membrane (22, 27). The broader and decomposable peak is caused by a variety of excited states and the diversity of the probe’s surroundings (microscopic viscosity, hydration degree, and so on) (28, 32). This complexity causes difficulty in analyzing Laurdan peaks. For GP, the selection of blue or red peaks are so variable that discrepancies can occur (32). The deconvolution analysis is effective, especially for separating the fluorescence emission spectra depending on the contribution of each of the excited states. In a recent study, the asymmetric function was selected to properly extract the red- and blue-shifted components (30). Though this asymmetrical deconvolution was consistent, double component deconvolution may not account for other molecular species.

Herein, the fluorescence emission spectra of Laurdan in PSM and DHPSM bilayers revealed the complexity of Laurdan’s location, with the broaden and decomposable emission peaks, confirming an observation in previous reports (12). In steady-state fluorescence measurements, the multiplicity of Laurdan’s location or interaction state in the membrane can be summarized in an emission spectrum. The broaden emission peaks are assumed due to the SM molecules forming intermolecular interactions and that Laurdan could have complex excited states in the lipid bilayers (12). The configurational restrictions and unique hydration properties in SM bilayers were confirmed based on the changes in anisotropy (Figure 2), the quick responses of the center of mass of the time-resolved emission spectra (Figure 3), and the blue shifts of the excitation spectra (Figure 4). The common GP analysis was also attempted. Figure S5 shows the transition in GP values due to increase in temperature in DPPC, DOPC, PSM, and DHPSM bilayers. Drastic shifts in GP values at around the phase transition were also found in DPPC, PSM, and DHPSM bilayers. However, considering the broaden emission peaks in SM bilayers (shown in Figure 1), the analysis based on GP values might have been concealing specific hydration properties in PSM and DHPSM bilayers. Based on the above results, information obtained from the Laurdan molecules is complex especially in SM bilayers and requires more careful interpretation. In the following section, the deconvolution of Laurdan spectrum was carried out, to investigate more appropriate hydration properties in SM bilayers.

### Deconvolution of the Laurdan spectra

In previous, a deconvolution of Laurdan spectrum into two components was employed in Laurdan studies, because the well-known emission peak at 440 nm and 490 nm correlated with the hydrated and less hydrated states, respectively (22). The specific Laurdan fluorescent properties of sphingolipids were confirmed through our results in this study, and it can be considered that Laurdan has more than two excited energy states. From the observed spectra in Figure 1, four components can be detected particularly for the Laurdan in SM bilayers, wherein each peak position at approximately 420 nm, 440 nm, 490 nm, and greater than 490 nm). Herein, the deconvolution into four components were performed.

Figure 5 shows the peak positions and the area ratios obtained from the four deconvolution curves. Based on the peak position changes to the increasing temperature, they can be classified into four groups (less than 440 nm, 440 ~ 460 nm, 460 ~ 480 nm, and greater than 480 nm). In DPPC, PSM, and DHPSM bilayers, the most blue-shifted peak at approximately 420 nm had disappeared during the phase transition. As suggested from the excitation spectra, this peak is originated from stabilizing effects, i.e., tight-packing or hydrogen bond interactions could prevent the solvation to fluorophore. Interestingly, in the loosely-packed bilayer (like DOPC), Laurdan also had a peak at 420 nm throughout the whole temperature range. This result seems contradictory, however, the Laurdan in DOPC bilayer had a short lifetime (~ 4 ns) at 420 nm of emission compared with that in SM bilayers and DPPC bilayer (~ 8 ns) (data not shown), suggesting that the blue shifted components can be derived from the collisional effects which could bring the excited Laurdan back to the ground state through the non-radiative pathway. During this pathway, most of the excited Laurdan can be quenched or relaxed.

**Figure 5.**
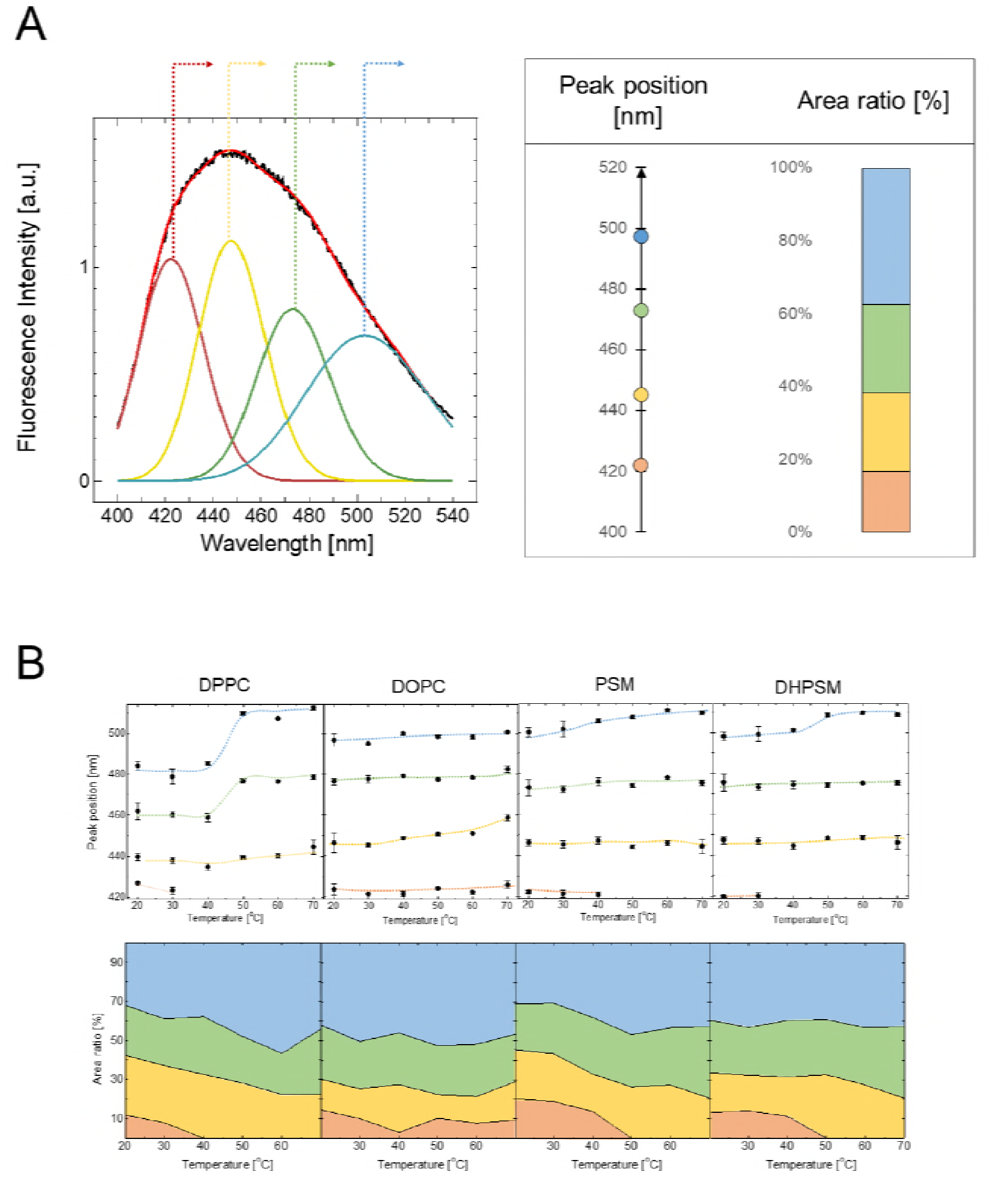
(A) Emission spectra of Laurdan in DPPC, DOPC, PSM, and DHPSM were deconvoluted using the lognormal fitting function as described in Eq. 3. From the four deconvolution curves, peak positions of each curve and the integrated area ratios to the whole spectral area were calculated. The peak positions and the integrated area percentages obtained for each bilayer were pictured in the corresponding color with the deconvolution spectra. (B; above) The peak positions of each deconvolution curve as a function of temperature, in DPPC, DOPC, PSM, and DHPSM bilayers. (B; below) The transition of area ratios of each deconvolution curve as a function of temperature, in DPPC, DOPC, PSM, and DHPSM bilayers.

Compared with the case in DPPC bilayer, the deconvolution curves in PSM and DHPSM bilayers showed more red-shifted peaks similar to the result in DOPC bilayer. In DPPC bilayer, the shift of peak position among the phase transition was observed; however, in SM bilayers, only slight red-shifts had been observed. These results indicate that the Laurdan molecules in SM bilayers can be distributing heterogeneously across the vertical direction of bilayer; one can be positioned into significantly hydrated region similar to the distribution in DOPC bilayer, and the other can be replaced in highly dehydrated region as showing the deconvoluted peak component at 420 nm.

For the interpretation of the deconvolution results, the number of peaks and their peak position can be linked to the hydration property of water molecules existing in bilayer interface. Parasassi *et al*. have previously evaluated the number and probability of water molecules that contribute to relaxation of Laurdan molecules using Poisson distribution (28): they found out that there would be between 0 and 5 water molecules. The three major hydration states with 1 to 3 water molecules could be more probable as compared with other states (0 or more than 4 water molecules). In this study, we also observed the four peak distributions in the time-resolved emission spectrum (less than 440 nm, 440 ~ 460 nm, 460 ~ 480 nm, and greater than 480 nm) which correspond to the deconvoluted peak distributions obtained from the results in Figure 5 (data not shown). Therefore, the four-component deconvolution approach is suitable, because Laurdan can take multiple excited states due to conformation restrictions caused by hydrogen bonding and the three measured relaxed states affected by the surrounding environment.

### Hydrogen bonding network and interfacial hydration states in SMs

According to previous studies of PSM and DHPSM bilayers, DHPSM bilayers have stronger intermolecular hydrogen network than PSM, which can be one of possible reasons to explain the differences in their physicochemical properties in bilayer (T_m_, PSM; 41.2 °C < T_m_, DHPSM; 47.7 °C) (39, 44). The immiscibility of dihydro-stearoyl-SM and DOPC bilayers was reported by Kinoshita *et al*., and this can be interpreted to be due to increased intermolecular hydrogen bonding (45). Regarding the anisotropy results and deconvolution dependencies on temperature (Figure 2 and 5, and Figure S5), the DHPSM bilayer showed a phase transition at higher temperatures than PSM bilayer, and the analyzed hydration properties of DHPSM bilayer was rather hydrophilic as compared to PSM bilayer. However, considering the effect of intermolecular hydrogen bonds, Laurdan emission spectra in PSM bilayers were largely blue-shifted compared to that in DHPSM bilayers (Figure 1). Furthermore, the Laurdan emission deconvolution analysis of PSM bilayers also revealed a relatively large contribution from the blue-shifted component (about 7% larger than DHPSM) (Figure 5). Regarding the extent of hydration, the DHPSM bilayer was more flexible and hydrated than that of PSM as suggested by the Laurdan emission spectra results, anisotropy and the center of mass of the time-resolved emission spectra (Figure 1, 2, and 3). The predominant blue-shift of emission spectra of Laurdan in PSM bilayer, and the highly hydrated properties in DHPSM bilayers seem contradictory to the previous observation of a strong intermolecular hydrogen bonding network found in DHPSM bilayer.

Yasuda *et al*., has also suggested that DHPSM has a more flexible head group compared to PSM based on quantum chemistry approach (46). They found that the properties of the 3OH group of DHPSM is different to those in PSM, because the *trans* double bond in PSM could restrict the rotational motion of C-C bond where 3OH group is present. The intramolecular hydrogen bond between 3OH group and phosphate oxygen is stronger in PSM than in DHPSM bilayer, as previously reported (39). It is considered that the restriction of 3OH group in PSM influences strong intramolecular hydrogen bonds, and the flexible 3OH group in DHPSM has more possibilities to form hydrogen bonds with other functional group. Besides, 2NH group of DHPSM could have hydrogen bonds with water molecules (39). Therefore, it is assumed that DHPSM has a superior accessibility to form hydrogen bond than PSM, which can stabilize the water molecules within the cavity around the flexible lipid head group. From the previous findings, the water exposure of lipids increases in the following order: PSM < DPPC < DHPSM, as measured from the fluorescence lifetime values of dansyl-PE under collisional quenching effect from D_2_O (12). This can explain the highly hydrated state of DHPSM and further suggests the existence of an interfacial hydration layer.

### Plausible model of hydration properties in PC and SM bilayers

A brief model for the membrane hydration and different populations of Laurdan according to fluorescence properties is illustrated in Figure 6. In DPPC bilayers, the Laurdan fluorescence indicated less hydration and high anisotropy below the phase transition (T < T_m_), and a shift to a higher hydrated state and a decrease of anisotropy above the T_m_. In DOPC bilayers, the membrane state is liquid-crystalline phase and acyl chains are in highly-disordered state. In PSM bilayers, some of Laurdan inserted into the membrane was strongly stabilized in the acyl chain region (“420” written with red font in Fig. 6). However, others can exist at the surface and sense different hydration states similar to those in DOPC bilayers. A similar situation could occur in DHPSM bilayers, but the significant difference is that DHPSM bilayers may retain more water molecules due to less intramolecular hydrogen bonds, leading to a high density of water molecules and stronger relaxation of Laurdan compared to that in PSM bilayers. Generally, the dipole moment of the lipid head group derived from zwitterionic part of the phosphocholine is aligned almost in parallel to the horizontal direction of the bilayer, in which the cationic choline part interacts with the anionic phosphate part of the neighboring lipid (40, 47, 48). Although this is just a speculation, the head group orientation could be restricted in SM bilayers in a direction more perpendicular to the bilayer surface due to the restriction of intramolecular hydrogen bonds between 3OH and phosphate oxygen. Particularly, DHPSM could have water molecules between lipid head groups, hence, its steric effect promotes the alignment of head group dipole moment.

**Figure 6.**
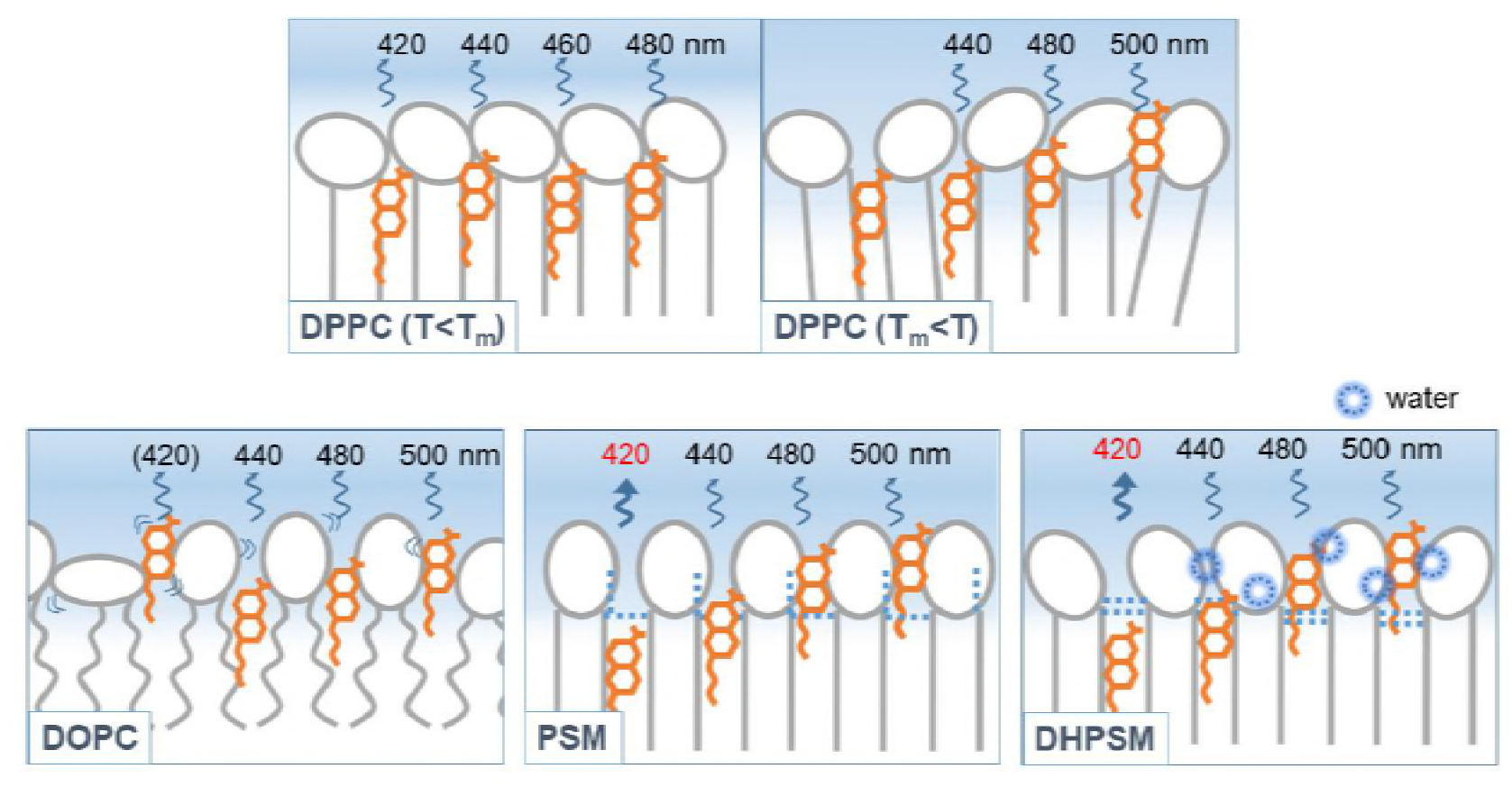
Hydration and the molecular model relation between lipid and Laurdan in DPPC, DOPC, PSM, and DHPSM bilayers. Lipids and Laurdan are illustrated in the color of gray and orange, respectively. The number written in the figure represent the emission peaks for each Laurdan molecule. “(420)” in DOPC is the component with high mobility and short lifetime. Red font of “420” is suggests that the component is stabilized by the hydrogen bonding network.

### Hydration water layer existing at the sphingolipid bilayer interface

The high-dense hydration water in SM bilayers, suggested in the above, seems to be contradictory, in comparison to previous studies. In molecular dynamics simulations, the water penetration across the bilayer was higher in DPPC than in PSM since stronger intermolecular hydrogen bonds in PSM expelled the water from the membrane (20). Investigation of fluidity and hydration states of raft system indicated that raft-forming bilayer (SM/Cholesterol = 65/35) were less hydrated than bilayer in the gel phase (DPPC) even though their fluidities values were similar (49). This discrepancy may be due to that the Laurdan in our study sensed the water molecules at the surface of the membrane near the bulk region. M’Baye *et al*., suggested that the vertical orientation of Laurdan could differ depending on the lipid species (49). SM bilayers can form highly ordered bilayer structures with intermolecular hydrogen bonds, hence, Laurdan could exist relatively close to the surface of the bilayer.

The water molecules at the surface of bilayer could be regarded as a hydration layer. Pal *et al*. suggest that the surface charge densities of the lipid head groups can modulate the dynamics of water molecules (50). In some discussions of computational simulations, lipid alignment was one of the possible factors that could stabilize or perturb the water-water hydrogen network (51). Furthermore, a bilayer composed by lipids with small head groups, such as 1-palmytoyl-2-palmytoyl-*sn*-glycero-3-phosphoethanolamine (DPPE), exhibited a stronger interfacial water network than other lipid species such as PCs. This also supports the correlation between the surface alignment of lipid bilayer and the stability of the hydration layer (18).

As discussed in the above, the specific intermolecular hydrogen bonds of sphingolipids may stabilize the alignment of the lipid head group, which finally results in the formation the hydration layer on the membrane surface. By deconvoluting the steady-state Laurdan emission spectrum, the hidden components, such as the Laurdan which might be stabilized via hydrogen bonding or that locating at more hydrophilic environment, can be shed a light. Regarding the thermodynamic properties for the pre and main transition of lipid bilayers studied using DSC, PSM and DHPSM showed smaller enthalpy change in pre transition (0.59 and 1.80 kJ/mol, respectively) compared to that of DPPC (3.39 kJ/mol). The surface topography in lipid bilayer differs before and after the transition (52). This slight enthalpy change observed in SM bilayers may indicate that the surface property was maintained by the fully hydrated lipid head groups during the pretransition. According to the dielectric dispersion analysis, the cooperativity of water molecules associating lipid bilayer was dominant in DHPSM than in DPPC, indicating the possibility to possess not only water molecules associating inter lipid region but also the one existing at lipid bilayer surface (See supporting information Figure S6 and S7). These suggestions are feasible studies for hydration water layer, and it is possible to consider that the retained water molecules could form the hydration layer on the DHPSM bilayer surface. As revealed based on the Laurdan studies, the DHPSM bilayers were in highly hydrated states, in which lipid molecules are flexible that allows a penetration of the surrounding water molecules locating on the membrane surface, which may result in the formation of hydration water layer. Though it is a hypothesis, the unique approach in this study will provide a novel insight into the hydration property of DHPSM bilayers, which leads us to much deeper understanding towards the lipid intermolecular interactions and the membrane surface hydration details.

## Author Contributions

N.W., K.S., J.P.S. and H.U. designed the research; T.N. and J.P.S. provided the analytical tools; N.W., Y.G., K.S. and T.N. performed the experiments and analyzed the data; N.W., K.S., T.N., J.P.S., and H.U. wrote the article.

## Acknowledgements

This work was primarily supported by the Japan Society for the Promotion of Science (JSPS) KAKENHI Grant-in-Aids for Scientific Research (A) (26249116), Grant-in-Aids for Young Scientists (B) (16K18279), Grant-in-Aids for Challenging Exploratory Research (T15K142040). N.W. expresses her gratitude for JSPS Research Fellow (JP18J11666) and Tobitate! (Leap for Tomorrow) Young Ambassador Program. The Slotte laboratory was supported by grants from the Sigrid Juselius, the Jane and Aatos Erkko, and the Magnus Ehrnrooth Foundations.

## Supporting Citations

References (53–55) appear in the Supporting Material.

## Graphical Abstract

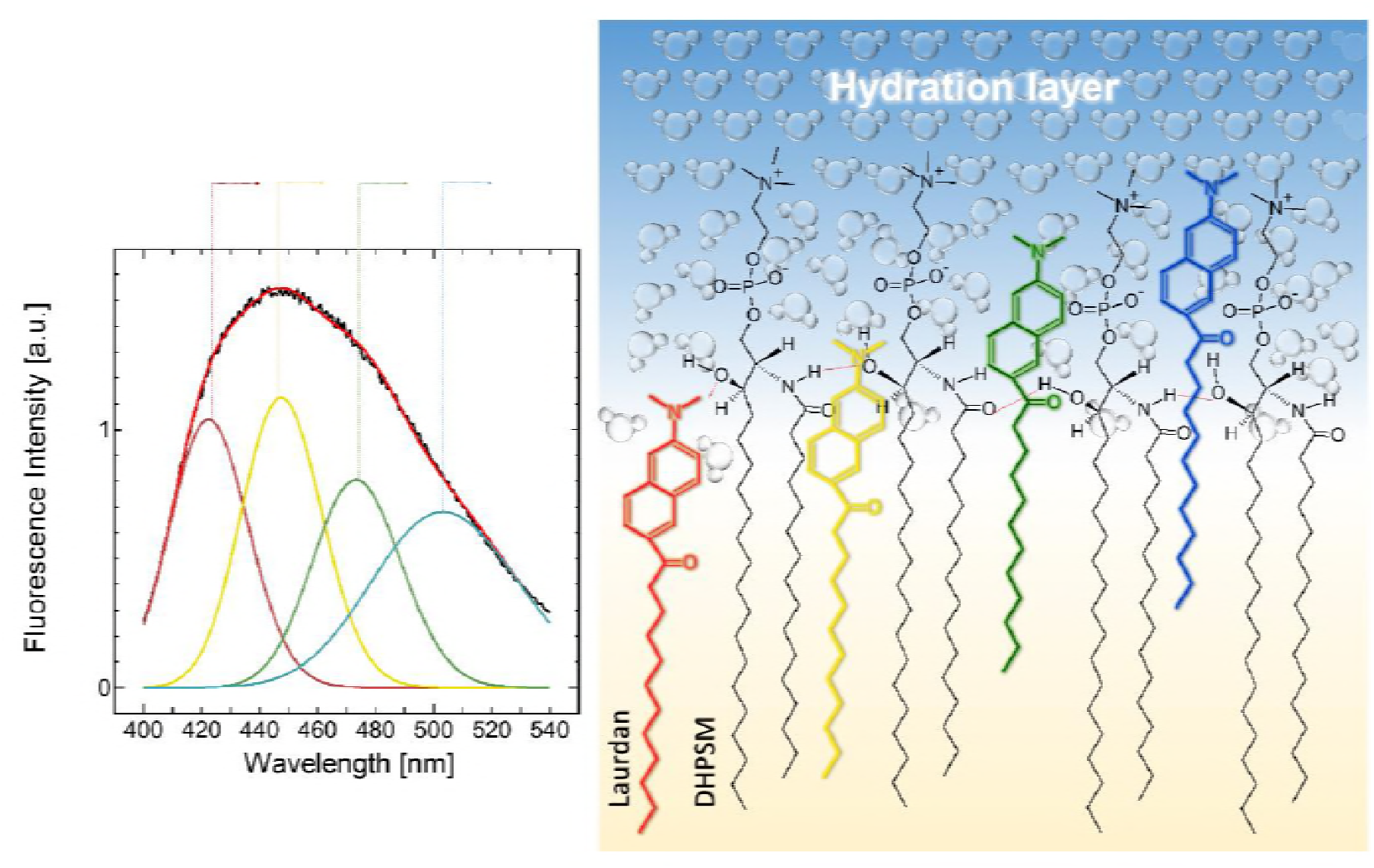

## References

1. Luckey, M. 2012. Membrane structural biology with biochemical and biophysical foundations. Cambridge: CambridgeUniversity Press.

2. Simons, K., and E. Ikonen. 1997. Functional rafts in cell membranes. Nature. 387: 569–572.

3. Barenholz, Y. 2004. Sphingomyelin and cholesterol: from membrane biophysics and rafts to potential medical applications. In: Membrane Dynamics and Domains. Springer, Boston, MA.

4. Casem, M.L. 2016. Membranes and membrane transport. In: Casem ML, editor. Case Studies in Cell Biology. Boston: Academic Press.

5. Lajoie, P., and I.R. Nabi. 2010. Lipid rafts, caveolae, and their endocytosis. In: International Review of Cell and Molecular Biology. Elsevier.

6. Slotte, J.P. 2016. The importance of hydrogen bonding in sphingomyelin’s membrane interactions with co-lipids. BBA. 1858: 304–310.

7. Ramstedt, B., and J.P. Slotte. Membrane properties of sphingomyelins. FEBS Lett. 531: 33–37.

8. Venable, R.M., A.J. Sodt, B. Rogaski, H. Rui, E. Hatcher, A.D. MacKerell Jr., R.W. Pastor, and J.B. Klauda. 2015. CHARMM all -atom additive force field for sphingomyelin: elucidation of hydrogen bonding and os positive cuvature. Biophys. J. 107: 134–145.

9. Ramstedt, B., and J.P. Slotte. 1999. Interaction of cholesterol with sphingomyelins and acyl-chain-matched phosphatidylcholines: a comparative study of the effect of the chain length. Biophys. J. 76: 908–915.

10. Kuikka, M., B. Ramstedt, H. Ohvo-Rekila, J. Tuuf, and J.P. Slotte. 2001. Membrane properties of D-erythro-N-acyl Sphingomyelins and their corresponding dihydro species. Biophys. J. 80: 2327–2337.

11. Epand, R.M. 2003. Cholesterol in bilayers of sphingomyelin or dihydrosphingomyelin at concentrations found in ocular lens membranes. Biophys. J. 84: 3102–3110.

12. Nyholm, T., M. Nylund, A.S. Söderholm, and J.P. Slotte. 2003. Properties of palmitoyl phosphatidylcholine, sphingomyelin, and dihydrosphingomyelin bilayer membranes as reported by different fluorescent reporter molecules. Biophys. J. 84: 987–997.

13. Nyholm, T., M. Nylund, and J.P. Slotte. 2003. A calorimetric study of binary mixtures of dihydrosphingomyelin and sterols, sphingomyelin, or phosphatidylcholine. Biophys J. 84: 3138–3146.

14. Alberts, B., A. Johnson, J. Lewis, D. Morgan, M. Raff, K. Roberts, and P. Walter. 2014. Molecular biology of the cell. 6th ed. New York and Abingdon, UK: Garland Science.

15. Jin, Z.-X., C.-R. Huang, L. Dong, S. Goda, T. Kawanami, T. Sawaki, T. Sakai, X.-P. Tong, Y. Masaki, T. Fukushima, M. Tanaka, T. Mimori, H. Tojo, E.T. Bloom, T. Okazaki, and H. Umehara. 2008. Impaired TCR signaling through dysfunction of lipid rafts in sphingomyelin synthase 1 (SMS1)-knockdown T cells. Int Immunol. 20: 1427–1437.

16. Villar, V.A.M., S. Cuevas, X. Zheng, and P.A. Jose. 2016. Localization and signaling of GPCRs in lipid rafts. In: Methods in Cell Biology. Elsevier. pp. 3–23.

17. Costard, R., I.A. Heisler, and T. Elsaesser. 2014. Structural dynamics of hydrated phospholipid surfaces probed by ultrafast 2D spectroscopy of phosphate vibrations. J. Phys. Chem. Lett. 5: 506–511.

18. Damodaran, S. 1998. Water activity at interfaces and its role in regulation of interfacial enzymes: a hypothesis. Colloids Surf., B. 11: 231–237.

19. Alarcón, L.M., M. de los Angeles Frías, M.A. Morini, M. Belén Sierra, G.A. Appignanesi, and E. Anibal Disalvo. 2016. Water populations in restricted environments of lipid membrane interphases. Eur. Phys. J. E. 39.

20. Saito, H., and W. Shinoda. 2011. Cholesterol effect on water permeability through DPPC and PSM lipid bilayers: a molecular dynamics study. J. Phys. Chem. B. 115: 15241–15250.

21. Sparr, E., and H. Wennerström. 2001. Responding phospholipid membranes—interplay between hydration and permeability. Biophys. J. 81: 1014–1028.

22. Parasassi, T., G. De Stasio, G. Ravagnan, R.M. Rusch, and E. Gratton. 1991. Quantitation of lipid phases in phospholipid vesicles by the generalized polarization of Laurdan fluorescence. Biophys. J. 60: 179–189.

23. De Vequi-Suplicy, C.C., C.R. Benatti, and M.T. Lamy. 2006. Laurdan in fluid bilayers: position and structural sensitivity. J. Fluoresc. 16: 431–439.

24. Bagatolli, L.A., T. Parasassi, G.D. Fidelio, and E. Gratton. 1999. A model for the interaction of 6-lauroyl-2-(n,n-dimethylamino)naphthalene with lipid environments: implications for spectral properties. Photochem. Photobiol. 70: 557–564.

25. Parasassi, T., E.K. Krasnowska, L. Bagatolli, and E. Gratton. 1998. Laurdan and prodan as polarity-sensitive fluorescent membrane probes. J. Fluoresc. 8: 365–373.

26. Vequi-Suplicy, C.C., K. Coutinho, and M.T. Lamy. 2014. Electric dipole moments of the fluorescent probes Prodan and Laurdan: experimental and theoretical evaluations. Biophys. Rev. 6: 63–74.

27. Parasassi, T., G. De Stasio, A. d’Ubaldo, and E. Gratton. 1990. Phase fluctuation in phospholipid membranes revealed by Laurdan fluorescence. Biophys. J. 57: 1179–1186.

28. Parasassi, T., E. Gratton, W.M. Yu, P. Wilson, and M. Levi. 1997. Two-photon fluorescence microscopy of laurdan generalized polarization domains in model and natural membranes. Biophys. J. 72: 2413–2429.

29. Harris, F.M., K.B. Best, and J.D. Bell. 2002. Use of laurdan fluorescence intensity and polarization to distinguish between changes in membrane fluidity and phospholipid order. BBA. 1565: 123–128.

30. Bacalum, M., B. Zorilă, and M. Radu. 2013. Fluorescence spectra decomposition by asymmetric functions: Laurdan spectrum revisited. Analytical Biochemistry. 440: 123–129.

31. Malacrida, L., S. Astrada, A. Briva, M. Bollati-Fogolín, E. Gratton, and L.A. Bagatolli. 2016. Spectral phasor analysis of LAURDAN fluorescence in live A549 lung cells to study the hydration and time evolution of intracellular lamellar body-like structures. BBA. 1858: 2625–2635.

32. Jay, A.G., and J.A. Hamilton. 2017. Disorder amidst membrane order: standardizing laurdan generalized polarization and membrane fluidity terms. J. Fluoresc. 27: 243–249.

33. Massey, J.B. 2001. Interaction of ceramides with phosphatidylcholine, sphingomyelin and sphingomyelin/cholesterol bilayers. Biochimica et Biophysica Acta (BBA) - Biomembranes. 1510: 167–184.

34. Hope, M.J., M.B. Bally, G. Webb, and P.R. Cullis. 1985. Production of large unilamellar vesicles by a rapid extrusion procedure. Characterization of size distribution, trapped volume and ability to maintain a membrane potential. BBA. 812: 55–65.

35. Rouser, G., S. Fleischer, and A. Yamamoto. 1970. Two dimensional thin layer chromatographic separation of polar lipids and determination of phospholipids by phosphorus analysis of spots. Lipids. 5: 494–496.

36. Haugland, R.P. 1996. Handbook of fluorescent probes and research chemicals. 6th ed. Eugene, OR: Molecular Probes Inc.

37. Lakowicz, J.R. 1999. Principles of fluorescence spectroscopy. New York: Kluvert Academic/Plenum Publishers.

38. Mohana-Borges, R., J. Lima Silva, and G. de Prat-Gay. 1999. Protein folding in the absence of chemical denaturants: Reversible pressure denaturation of the noncovalent complex formed by the association of two protein fragments. J. Biol. Chem. 274: 7732–7740.

39. Talbott, C.M., I. Vorobyov, D. Borchman, K.G. Taylor, D.B. DuPré, and M.C. Yappert. 2000. Conformational studies of sphingolipids by NMR spectroscopy. II. Sphingomyelin. BBA. 1467: 326–337.

40. Gally, H.U., W. Niederberger, and J. Seelig. 1975. Conformation and motion of the choline head group in bilayers of dipalmitoyl-3-sn-phosphatidylcholine. Biochemistry. 14: 3647–3652.

41. Frank, J. 1926. Elementary processes of photochemical reactions. Transactions of the Faraday Society. 21: 536–542.

44. Ferguson-Yankey, S.R., D. Borchman, K.G. Taylor, D.B. DuPré, and M.C. Yappert. 2000. Conformational studies of sphingolipids by NMR spectroscopy. I. Dihydrosphingomyelin. Biomembranes. 1467: 307–325.

45. Kinoshita, M., N. Matsumori, and M. Murata. 2014. Coexistence of two liquid crystalline phases in dihydrosphingomyelin and dioleoylphosphatidylcholine binary mixtures. BBA. 1838: 1372–1381.

46. Yasuda, T., M.A. Al Sazzad, N.Z. Jäntti, O.T. Pentikäinen, and J.P. Slotte. 2016. The influence of hydrogen bonding on sphingomyelin/colipid interactions in bilayer membranes. Biophys. J. 110: 431–440.

47. Griffin, R.G., L. Powers, and P.S. Pershan. 1978. Head-group conformation in phospholipids: a Phosphorus-31 nuclear magnetic resonance study of oriented monodomain dipalmitoylphosphatidylcholine bilayers. Biochemistry. 17: 2718–2722.

48. Büldt, G., H.U. Gally, and J. Seelig. 1979. Neutron diffraction studdies on phosphatidylcholine model membranes I. Head group conformation. J. Mol. Biol. 134: 673–691.

49. M’Baye, G., Y. Mély, G. Duportail, and A.S. Klymchenko. 2008. Liquid ordered and gel phases of lipid bilayers: fluorescent probes reveal close fluidity but different hydration. Biophys. J. 95: 1217–1225.

50. Pal, S., N. Samanta, D. Das Mahanta, R.K. Mitra, and A. Chattopadhyay. 2018. Effect of phospholipid headgroup charge on the structure and dynamics of water at the membrane interface: a terahertz spectroscopic study. J. Phys. Chem. B. 122: 5066–5074.

51. Tieleman, D.P., S.J. Marrink, and H.J.C. Berendsen. 1997. A computer perspective of membranes: molecular dynamics studies of lipid bilayer systems. BBA. 1331: 235–270.

52. Heimburg, T. 2000. A model for the lipid pretransition: coupling of ripple formation with the chain-melting transition. Biophys. J. 78: 1154–1165.

53. George, D.K., A. Charkhesht, O.A. Hull, A. Mishra, D.G.S. Capelluto, K.R. Mitchell-Koch, and N.Q. Vinh. 2016. New insights into the dynamics of zwitterionic micelles and their hydration waters by gigahertz-to-terahertz dielectric spectroscopy. J. Phys. Chem. B. 120: 10757–10767.

54. Hatakeyama, H., H. Akita, and H. Harashima. 2011. A multifunctional envelope type nano device (MEND) for gene delivery to tumours based on the EPR effect: A strategy for overcoming the PEG dilemma. Adv. Drug Deliv. Rev. 63: 152–160.

55. Disalvo, E.A., and M.A. Frias. 2013. Water State and Carbonyl Distribution Populations in Confined Regions of Lipid Bilayers Observed by FTIR Spectroscopy. Langmuir. 29: 6969–6974.

